# Brain Network Dynamics from Neuronal Avalanches for Longitudinal Motor-Imagery Decoding

**DOI:** 10.64898/2026.11.09.765302

**Authors:** Camilla Mannino, Alice Longhena, Mario Chavez, Marie-Constance Corsi

## Abstract

Brain–computer interfaces (BCIs) offer opportunities for personalized neurorehabilitation, but inter-session variability, repeated calibration, and the need for expert supervision limit their use beyond supervised settings. We propose a longitudinal BCI framework that integrates neuronal avalanches dynamics, reflecting cascades of propagation neural activity, with graph-theory and data-driven region of interest analysis to extract computationally efficient and neuro-physiological interpretable features. Source-reconstructed EEG from 20 BCI-naïve participants performing right-hand motor imagery (MI) and rest across four training sessions was represented by a directed weighted connectivity network based on neuronal avalanches. Local, mesoscopic, and global network measures revealed task-related changes predominantly in contralateral sensorimotor regions and distributed reorganization across training. We modelled here the learning-related evolution of feature distributions across previous sessions and used this trajectory to predict class-conditional distributions in an unseen fourth session, anticipating future brain states before new-session calibration. PageRank-based ROI selection achieved an AUC of 73.5 ± 7.9%, compared with 64.5 ± 10.6% using all ROIs, while reducing computational time by approximately 39% relative to performance-driven selection. The proposed framework provides interpretable, compact descriptions of avalanche propagation and exploits training-related changes for prospective decoding. Moving spatial selection and model optimization to an initial offline phase may reduce subsequent calibration demands and facilitate more autonomous BCI use.

## 1. INTRODUCTION

Brain–computer interfaces (BCIs) translate brain activity into commands for external devices and can support communication, motor control, and neurorehabilitation[1]. However, the daily-life use of non-invasive EEG-based BCIs remains constrained by low signal-to-noise ratio, long time preparation, repeated calibration, and experts supervision[2]. Inter- and intra-subject variability further influence both BCI learning and decoding performance[3], [4]. Features capable of identifying the current mental state while also capturing training-related adaptation could therefore support more individualized and efficient BCI use. Conventional EEG pipelines typically rely on spatially localized representations, such as sensorimotor rhythms or spectral-power modulations. Although these approaches can achieve high decoding performance[5], they often provide limited insight into the neural mechanisms underlying BCI control. Consequently, increasing attention has shifted from isolated regional activity toward interactions among distributed brain areas. Functional connectivity quantifies statistical relationship among brain regions signals over time, while graph-theoretical measures characterize the resulting brain networks at local, mesoscopic and global scales [6]. Compared with standard functional connectivity matrices (covariance or synchrony matrices), connectivity graphs provide a more interpretable representation of brain dynamics, with potential benefits for computational efficiency in BCI applications[7], [8].

Growing evidence indicates that graph-theoretical measures can serve both as markers of BCI-related brain reorganization and as features for decoding mental states[7]. Motor learning and BCI training are associated with changes in clustering, efficiency, integration, modularity, and connectivity degree, linking network topology to plasticity, behavioural improvement, and individual learning rate[9]-[10]. Connectivity measures such as node strength and centrality can also discriminate motor imagery from rest or between imagery classes, with multimeric approaches achieving performance comparable to standard parameters, such as power spectral density and common spatial patterns[11]-[12].

Neuronal avalanches provide a complementary representation by capturing the spatiotemporal propagation of both oscillatory and aperiodic activity across cortical regions[13], [14], [15]. Their propagation can be encoded through transition matrices describing the probability that activation in one region is followed by activation in another. Avalanche-based connectivity has already been shown to distinguish motor imagery from rest[16], [17], while avalanche length and activation count capture both task-related and longitudinal training effects and correlate with BCI performance[18]. These findings motivate the use of graph descriptors of avalanche propagation to jointly characterize motor-imagery state, training-related reorganization, and prospective decoding.

In this work, we investigate whether the topological organization of neuronal-avalanche propagation captures both the BCI-related brain state and its evolution across training, and whether this information can support prospective decoding within a reduced, physiologically meaningful feature space. We hypothesized that multiscale graph measures distinguish MI from rest, track training-related reorganization, and contain longitudinally predictive information that can support subject-specific classification, whereby the evolution of rest and MI dynamics across previous sessions can be used to discriminate brain states in a subsequent unseen session. We further tested whether task- and training-informative ROIs could be identified during calibration to preserve decoding performance while reducing dimensionality and computational cost. Overall, the framework evaluates whether feature and spatial optimization can be shifted partly to an offline calibration phase, enabling interpretable, lower-cost decoding in subsequent sessions and potentially supporting more efficient and autonomous BCI-based rehabilitation.

## 2. METHODS

### 2.1 Participants & Protocol

Twenty healthy, right-handed BCI-naïve adults (mean age = 27.5 ± 4.0 years; 12 men) without medical or psychological disorders participated in the study[19], [20]. The French CPP-IDF-VI ethics committee approved the protocol; participants provided written informed consent and received financial compensation. The study followed the Declaration of Helsinki. Four EEG-based BCI sessions were conducted over two weeks, with 10-minute home-training videos between laboratory sessions. Participants controlled a horizontally moving cursor’s vertical position through sensorimotor α and/or β activity: right-hand grasping imagery reached the upper target, and rest reached the lower target. MI and REST trials were equally represented and randomized. EEG was recorded from 74 channels at 1 kHz; T9 and T10 were removed and data downsampled to 250 Hz. Infomax ICA was applied to correct artifacts, with fewer than 10% of trials rejected. Individual BEM head models and weighted minimum-norm estimation supported source reconstruction. Activity was projected to MNI-ICBM152 and parcellated with the Desikan–Killiany atlas[21]. Further details on preprocessing and source reconstruction are provided in Corsi et al. (2020)[10].

### 2.2 Brain Network definition

Neuronal avalanches were identified from continuous EEG recordings by first z-scoring the time series of each brain region and subsequently converting them into binary activation patterns through amplitude thresholding. An avalanche began when at least one region exceeded the amplitude threshold θ_av_ and ended when all regions returned below threshold. Events shorter than a minimum duration λ_min_av_ were excluded as noise or artifacts. One predefined parameter combination (θ_av_, λ_min_av_) was used throughout, following the selection approach in [18], [22].

Once detected, neuronal avalanches were encoded using the weighted-stochastic Avalanche Transition Matrix (W), which provides a probabilistic representation of the spatiotemporal propagation of activity across brain regions. For each avalanche, directed transitions were defined between cortical regions active at consecutive time points within a row-substochastic formulation. To obtain a single representation of avalanche dynamics for each trial, the transition matrices derived from all avalanches detected within that trial were combined using a duration-weighted average. The resulting W can therefore be interpreted within a Markov-chain framework. A detailed description of the W formulation and construction is provided in [22].

Each trial-specific *W ∈* ℝ^*N*×*N*^ was mapped onto a directed, weighted brain network[23]. N corresponds to the cortical ROIs and each non-zero off-diagonal, W_i,j_, represents the transition probability from source node (i) to target node (j) and defines the weight of the corresponding directed edge. By construction, W is a substochastic matrix, such that the outgoing transition weights from each source node are normalized to sum to one while rows corresponding to nodes with no observed outgoing transitions remain zero.

Accordingly, while total outgoing connectivity is constrained by normalization, incoming connectivity remains free to vary across nodes according to the distribution of transition probabilities converging onto each region. Direction-dependent network metrics were therefore evaluated in the incoming direction whenever applicable. Self-connections (W_ii_) were excluded, and all positive transition probabilities were retained without additional thresholding. The resulting networks therefore captured the direction and relative probability of neuronal-avalanche propagation across cortical regions and were constructed independently for each trial, participant, experimental condition, and training session. The networks are generated using NetworkX toolbox in Python[24].

### 2.3 Brain Network Metrics

We characterize the topological organization of the reconstructed brain networks at different spatial scales: local, mesoscopic, and global[7], [23].

At *local-scale* we considered the node strength[25], communicability[26] and local efficiency[27] to characterize complementary aspects of the topological and communication role of individual cortical regions.

#### Local incoming node strength

The incoming strength of node j, 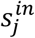, was defined as the cumulative weight of all directed transitions weights reaching that node[25]:

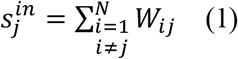

#### Local incoming communicability

Communicability[26] was used to quantify the accessibility of each node by accounting for both direct and indirect propagation pathways. For each node, j, incoming communicability was defined as:

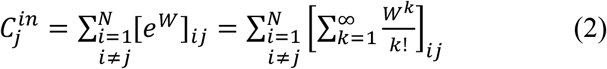

Where k denotes the length of a walk through the network, i.e., the number of successive transitions connecting two nodes. The term [*W*^*k*^]_*ij*_ represents the contribution of all walks of length k from node i to node j, whereas the factor (1/k!) progressively down-weights the contribution of longer walks. Thus, k=1 corresponds to direct transitions whereas k>1 accounts for indirect walks through the network.

#### Local efficiency

Local efficiency[27] was used to quantify the efficiency of communication among the neighborhood nodes of each region. For node j, local efficiency was defined as the efficiency of the subgraph G_j_ induced by the neighborhood of node j:

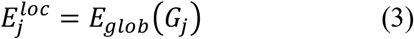

Nodal efficiency was computed using the weighted directed formulation implemented in the Brain Connectivity Toolbox.

At *global scale*, incoming node strength and communicability were obtained by averaging the corresponding local measures across all nodes (N)[28] to characterize the network as a whole and provide summary measures of its overall macro-scale properties.

**Global incoming node strength:**

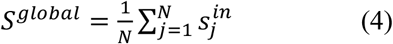

**Global incoming communicability:**

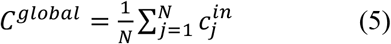

**Global efficiency:**

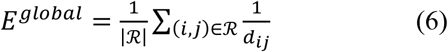

Where ℛ is the set of ordered node pairs connected by a finite directed path, and d_ij_ is the weighted directed shortest-path distance[29] from node i to node j.

At the *mesoscopic scale*, we characterized intermediate network organization using two complementary approaches: directed three-node motifs (triads)[30], capturing recurrent local interaction patterns, and core–periphery organization[31], describing the relative hierarchical position of nodes within the network.

#### Directed triads

Triadic motifs were used to identify recurrent patterns of directed interaction among small groups of cortical regions, providing a mesoscopic description of how local propagation pathways are organized beyond single-node properties[7]. Directed three-node interaction patterns[30] were quantified independently for each trial and experimental condition. For motif identification, the weighted transition matrix was converted into a binary directed adjacency matrix, A_ij_, such that the motif analysis described the presence and orientation of directed transitions rather than their magnitude. For each three-node subnetwork, the complete directed connectivity pattern was then encoded according to the six possible pairwise transitions. The different combinations of these directed transitions defined a set of 18 ordered triadic motifs (see Fig. 1a).

**Fig. 1:**
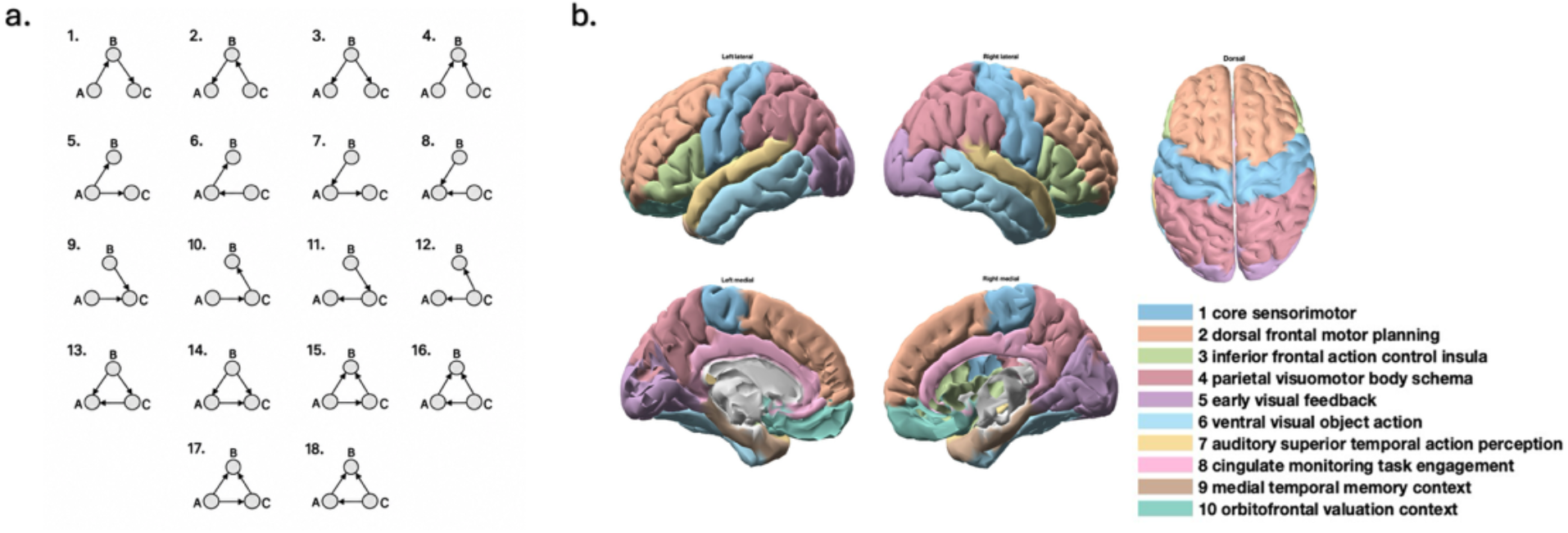
Overview of directed triad configurations and brain functional group parcellation. a) Schematic representation of the 18 possible directed triad configurations, illustrating the directionality of connections between three nodes (A, B, and C). b) Anatomical visualization of the ten functional groups defined according to the DK10 parcellation, displayed across lateral, dorsal, and medial views of the brain hemispheres. Each colour represents a distinct functional group.

To obtain a lower-dimensional and functionally interpretable representation of triadic organization, triads were regrouped post hoc according to the functional roles of their constituent cortical regions. This reassignment did not involve constructing a separate network with functional groups as nodes. Cortical regions were assigned to 10 functional groups relevant to motor imagery, action planning, sensory processing, and visual feedback (Fig. 1b and Supplementary Table 1). Triads sharing both the same directed motif and the same ordered functional-group triplet were then aggregated. For each specific triad, its presence was assessed in each trial, and its occurrence across trials was subsequently quantified. This analysis was performed independently for each subject, session, and experimental condition, allowing triadic organization to be characterized separately during motor imagery and rest across training sessions.

#### Core-Periphery

This analysis was used to describe the hierarchical position of cortical regions within a more central core or a peripheral network structure. Core–periphery organization was characterized using the approach proposed in [31], adapted to the weighted directed nature of the transition networks. Nodes were ranked in descending order of incoming strength, with stronger incoming connectivity associated with lower ranking,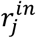:

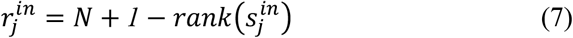

Where N is the number of nodes and 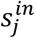 correspond to the incoming node strength.

All network metrics, except triads, were first estimated independently for each trial and subsequently averaged across the trials belonging to the same subject, session, and experimental condition. The same procedure was applied to the core–periphery analysis. This procedure yielded one node-specific value for each local metric, including incoming rank, and one network-level value for each global metric, while preserving trial-level variability during metric estimation.

### 2.4 Statistical Analysis of network effects

The statistical analysis aimed to assess task-related and training-related changes in network organization at the group level. Network measures were first computed at the individual-subject level, yielding one representative value for each subject, session, and experimental condition. These subject-specific values were then used as the observations in the group-level statistical analyses, with subjects constituting the statistical sample. This framework was applied across the different levels of network organization, including individual edge weights, node-level (local) network metrics, the occurrence of specific directed triads within predefined functional-group configurations, and the incoming rank derived from the core–periphery analysis. Task-related effects were assessed by comparing rest and MI. Within each session using paired Wilcoxon signed-rank tests[32] and/or using the t-test were performed across subjects, with Benjamini–Hochberg false-discovery-rate (FDR)[33] correction for multiple comparisons. In addition, a two-way rank ANOVA[34] was used to evaluate the overall main effect of condition across sessions. Training-related effects were assessed by evaluating changes across the four sessions. Friedman tests[35] where performed separately within rest and MI state, with FDR correction[33], while the main effect of session from the two-way rank ANOVA[34] was used to characterize the overall evolution of network properties across training sessions irrespective of condition.

### 2.5 Data-driven ROIs selection

To identify cortical regions whose network relevance changed jointly as a function of experimental condition and training, we implemented a data-driven ranking procedure. In networks theory, PageRank assigns each node an importance score according to its contribution importance in the network based on the weighted directed connections it receives[36]. Here, PageRank was computed directly from the W_ij_ using the Python scikit-network[37] implementation, yielding one value for each ROI and trial, separately for each subject, condition, and session. A subject-level statistical analysis was then performed. For each subject, the trial-wise PageRank values were entered, independently for each ROI, into a two-way ANOVA and selection was based specifically on Condition x Session interaction. This criterion identified regions in which the REST–MI difference in PageRank changed across training sessions, rather than regions showing only effect of condition or session. Interaction p-values were corrected across ROIs using the FDR procedure, and ROIs showing a significant FDR-corrected interaction were retained. Subject-specific ROI sets were then combined across participants, and only regions identified in most subjects were included in the final PageRank-based subset.

PageRank was chosen as an independent network-based selection criterion because it is naturally suited to the weighted, directed stochastic structure of W_ij_ and provides a spatial-selection measure distinct from the graph features subsequently used for classification. Its effectiveness was evaluated within the classification framework by comparing to other three ROI configurations: i) an all-ROI model without spatial selection; ii) an a-priori selection literature-informed subset comprising bilateral precentral, paracentral, postcentral, and caudal middle frontal regions together with the left superior frontal ROI, based on their established involvement in motor imagery and sensorimotor processing[38], [39]; and iii) an AUC-based data-driven strategy derived from cross-validation performance[40].

In classification framework the PageRank-based ROI selection was performed using exclusively data from the first three sessions, without incorporating any information from the fourth session. In this way, the fourth session was preserved as a completely unseen test set, thereby preventing information from the test data from influencing feature selection and reducing the risk of overfitting and data leakage.

### 2.6 Prediction and Classification Model

A subject-specific longitudinal prediction and classification framework was developed to determine whether the evolution of brain-network properties across BCI training could support single-trial discrimination between rest and motor imagery in a subsequent unseen session. Input features comprised node strength, communicability, and efficiency at both global and local scales, with local measures represented ROI-wise and evaluated using the alternative ROI configurations described in Section 2.4. Trial-wise feature values were log-transformed as *x*^*′*^ = *log*[*max*(*x, ε*)] with *ε* = *10*^*−12*^ prevented undefined values for zero observations, and their REST and MI distributions were modelled using Gaussian probability density functions (Fig. 2, steps 1–2).

**Fig. 2:**
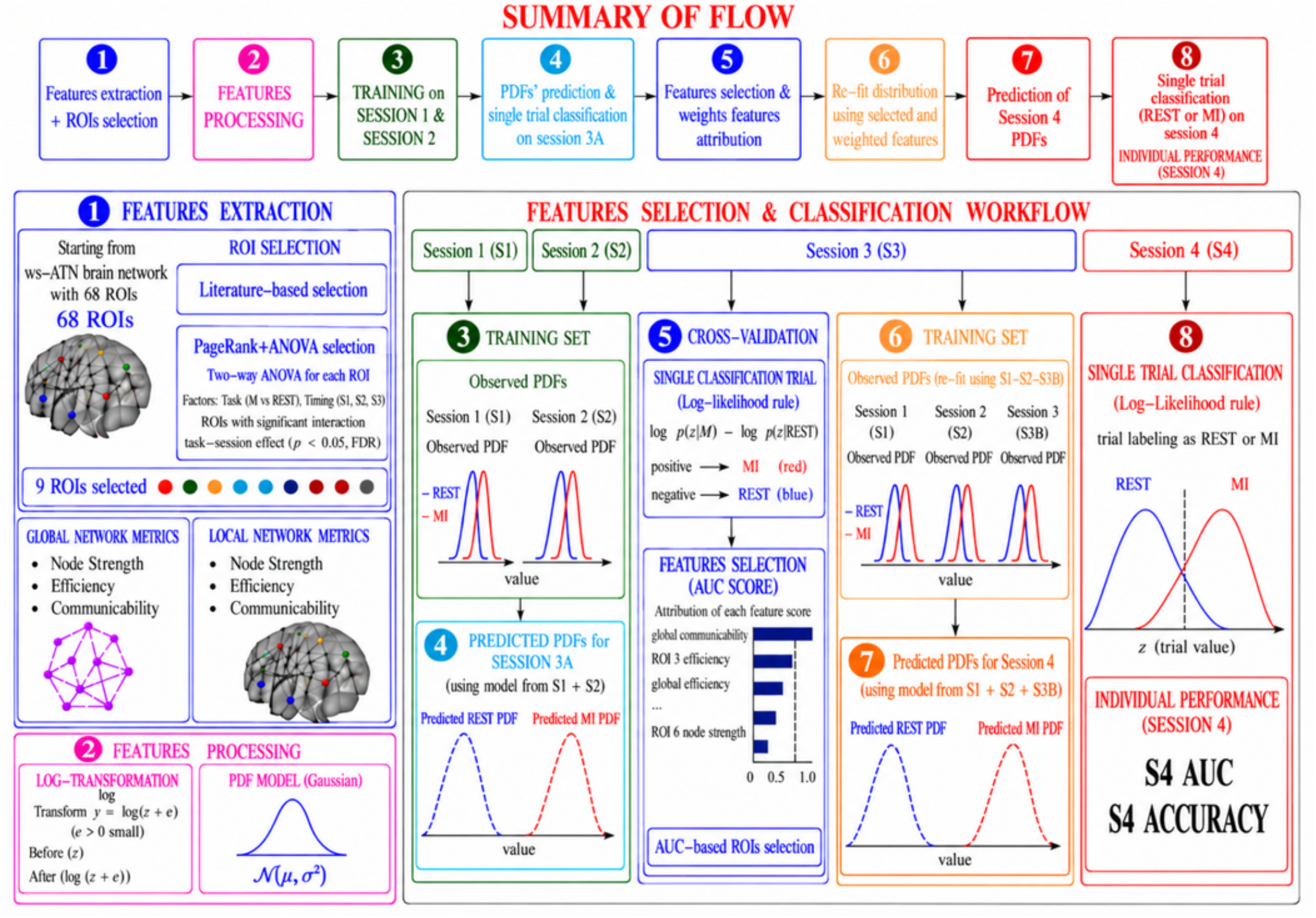
Overview of the feature selection, prediction, and classification workflow. The pipeline comprises eight steps: (1) extraction of global and local network metrics from selected ROIs; 2) feature preprocessing and Gaussian probability density function (PDF) modeling; 3) estimation of resting-state (REST) and motor imagery (MI) feature distributions using sessions S1–S2; 4) prediction of S3 distributions; 5) AUC-based feature selection and weighting through cross-validation; 6) refinement of feature distributions using S1–S3 data; 7) prediction of S4 distributions; and 8) single-trial REST/MI classification using a log-likelihood decision rule, with performance assessed by AUC and accuracy.

To preserve the longitudinal structure of the protocol and prevent information leakage, data were partitioned chronologically. Sessions 1 and 2 formed the initial training set, while Session 3 was divided into two non-overlapping halves: S3A was used exclusively as cross-validation set for parameters, features and strategy selection, and S3B was withheld for subsequent final model fitting. Session 4 was retained as an independent test set and was accessed only after all model parameters and selection criteria had been fixed. For each subject and network feature, predictions were performed according to four candidate temporal-evolution rules: pooled estimation, which assumes a stable underlying distribution across sessions[41]; recency-weighted estimation, which gives greater influence to the most recent observations[42]; linear-trend extrapolation, which projects the estimated temporal trend forward[43]; and extrapolation of the most recent session-to-session change, which assumes that the latest observed change persists into the next session[44]. Rather than selecting a single temporal rule, the resulting Gaussian predictions were combined into class-specific PDF mixtures using non-negative weights constrained to sum to one and optimized on S3A through a class-balanced logistic loss with L2 regularization (Fig. 2, steps 3–4).

Single-trial classification on S3A was based on the difference between the predicted class log-likelihoods:

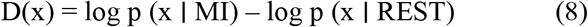

with positive values favouring MI and negative values favouring REST. Cross-validation was used to perform the features selection, choosing among global, local, and combined scales together.

For the AUC-based approach, each ROI was ranked according to the discriminative performance of its associated local network measures, namely node strength, efficiency, and communicability. The primary ranking criterion was the highest cross-validation AUC obtained among the three measures for that ROI, while the mean AUC across the measures was used only to resolve ties. Whenever an ROI was retained, all local network-feature components associated with that ROI were included in the corresponding fusion model.

Here, standardized feature scores were weighted according to their cross-validation AUC, with non-negative weights based on performance above chance level[45] and normalized to sum to one. For each subject, the final configuration was selected primarily according to cross-validation AUC, followed by balanced accuracy and accuracy, with the lower-dimensional solution preferred when performance was equivalent (Fig. 2, step 5).

After cross-validation, all model parameters and selection criteria were fixed. The model was refitted using Sessions 1, 2, and S3B to predict the REST and MI distributions of Session 4, which remained completely unseen during model development. Session 4 trials were classified using these predicted distributions and the decision threshold calibrated on S3A (Fig. 2, steps 6–8). Classification performance was quantified separately for each subject using ROC-AUC, accuracy, balanced accuracy, class-specific precision, recall, F1-score, and the corresponding confusion-matrix entries.

## 3. RESULT

### 3.1 Motor Imagery training induces brain network re-organization

We evaluated the spatiotemporal cortical changes associated with BCI training by investigating two distinct effects: the training-effect and the condition effect, through statistical analyses of different graph-theoretical metrics across subjects.

A significant training-effect was detected at both *the global level* (node strength: *p* = image; communicability: *p* = image; efficiency: *p* = 0.002; all values are reported after FDR correction; Supplementary Fig. 1) and the local level. This effect was largely driven by the motor imagery condition, as revealed by Friedman tests performed separately for each condition (Figure 3). This distinction was particularly evident *at the local level*. During REST, only the right caudal anterior cingulate cortex exhibited a significant session effect, restricted to node strength and communicability. In contrast, MI showed a more widespread pattern of session-dependent changes, involving sensorimotor, frontal, cingulate, parietal, and medial temporal regions, particularly in terms of communicability.

**Fig. 3:**
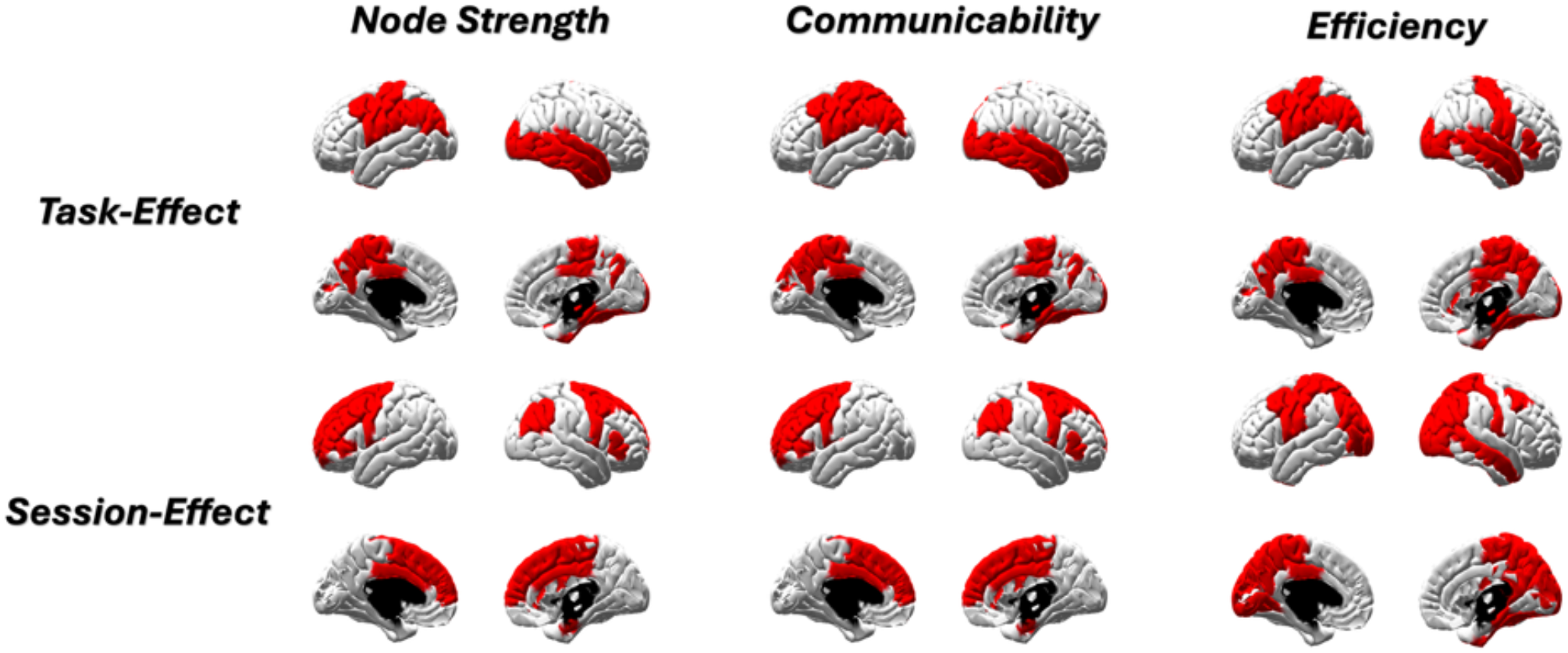
Spatial distribution of significant task and session effects across network metrics. ROIs showing significant effects (FDR-corrected p < 0.05) identified by two-way ANOVA for node strength, communicability, and efficiency are highlighted in red. The upper row shows the main effect of task (MI vs. REST) across all four sessions, while the lower row shows the main effect of session (S1–S4) across both conditions. Additional condition- and session-specific analyses are presented in Supplementary Fig. 2.

Overall, these findings suggest that training-related network reorganization was predominantly associated with motor imagery and extended beyond the sensorimotor system to encompass distributed frontal, cingulate, parietal, and associative regions.

Based on this evidence of distinct training-related responses between the two brain states, we further investigated the condition effect across sessions by comparing graph-theoretical measures between the MI and REST conditions across subjects within each individual session.

*At the global scale*, only efficiency showed a significant task effect in Sessions 3 and 4 (q = 0.036 and q = 0.033, respectively), with higher values during Rest (Supplementary Fig. 1) while *at the nodal level*, task-effect across the four training sessions, assessed using the two-way ANOVA, showed many significant ROIs predominantly left-lateralized (Fig. 3). Node strength and communicability were significantly modulated mainly in the left precentral, postcentral, paracentral, and precuneus regions, whereas local efficiency showed a broader distribution involving bilateral sensorimotor and parietal areas, posterior cingulate, precuneus, and additional frontal and fusiform regions. The left precentral, postcentral, paracentral, and precuneus cortices were consistently identified across all three nodal measures, indicating a robust involvement of controlateral-lateralized sensorimotor and parietal regions in the MI–Rest contrast. Within individual sessions, Wilcoxon signed-rank tests with FDR correction revealed a clear session-dependent evolution of the MI–Rest difference (Supplementary Fig. 2). In Session 1, significant effects were observed in the left postcentral cortex for node strength and communicability, while local efficiency additionally involved the left precentral cortex. No significant ROIs were detected in Session 2. In Session 3, communicability identified the left postcentral and supramarginal regions, whereas local efficiency showed a broader pattern involving left frontal, sensorimotor, cingulate, and parietal regions. In Session 4, a similar spatial organization was observed, with node strength also showing significant effects; notably, the left paracentral cortex was significant across node strength, communicability, and local efficiency. Across these significant nodal effects, values were generally higher during Rest than MI, indicating a reduction of these network properties during motor imagery. These network metrics results reflect in more comprehensive and interpretable way the same finding observed at edgewise network analysis. *At single scale edge*, after FDR correction, significant MI–Rest differences remained only in Sessions 1 and 4. Session 1 showed a sparse and focal pattern cantered mainly on the left postcentral cortex, whereas Session 4 exhibited a denser and more distributed network involving the left precentral, postcentral, supramarginal, and paracentral regions, with the left paracentral cortex emerging as a prominent hub with a large number of significant incident connections. The REST-MI contrast showed predominantly higher W_ij_ transition values during Rest than MI in left sensorimotor connections, whereas the opposite pattern was more frequently observed in right-hemisphere regions (Supplementary Fig. 3).

### 3.2 Brain network organization at mesoscopic scale

#### Directed Triads

Ten triads met the predefined selection criteria: a significant session effect during MI, a significant MI–REST difference in at least one session, and an average occurrence of at least 50% in at least one session. Considering the functional composition of these triads (Fig. 4a), the three most frequently represented functional groups were parietal visuomotor body schema, cingulate monitoring/task engagement, and core sensorimotor. Their distribution was also position-dependent within the triads. The cingulate monitoring/task-engagement group occurred predominantly at node position 1, the core sensorimotor group was most frequently represented at node position 2, whereas the parietal visuomotor body-schema group showed a particularly strong preference for node position 3. This positional organization suggests that the selected triads were characterized not only by recurrent combinations of functional systems, but also by a preferential arrangement of these systems within the local connectivity motifs. For all ten selected triads, a significant task-related difference between MI and REST was observed in the final training session (S4) (Fig. 4b). In every case, triad occurrence was higher during motor imagery than during rest, indicating a consistent MI-related enhancement of these specific connectivity patterns at the end of training. At the same time, the subject-averaged occurrence of the selected triads showed an overall decreasing trend across training sessions in both MI and REST conditions, although the magnitude of this decrease differed across triads (Fig. 4b). Together, these results suggest that training was associated with a progressive reduction in the overall occurrence of these local connectivity configurations, while the relative distinction between MI and REST remained evident and became consistently expressed in the final training session.

**Fig. 4:**
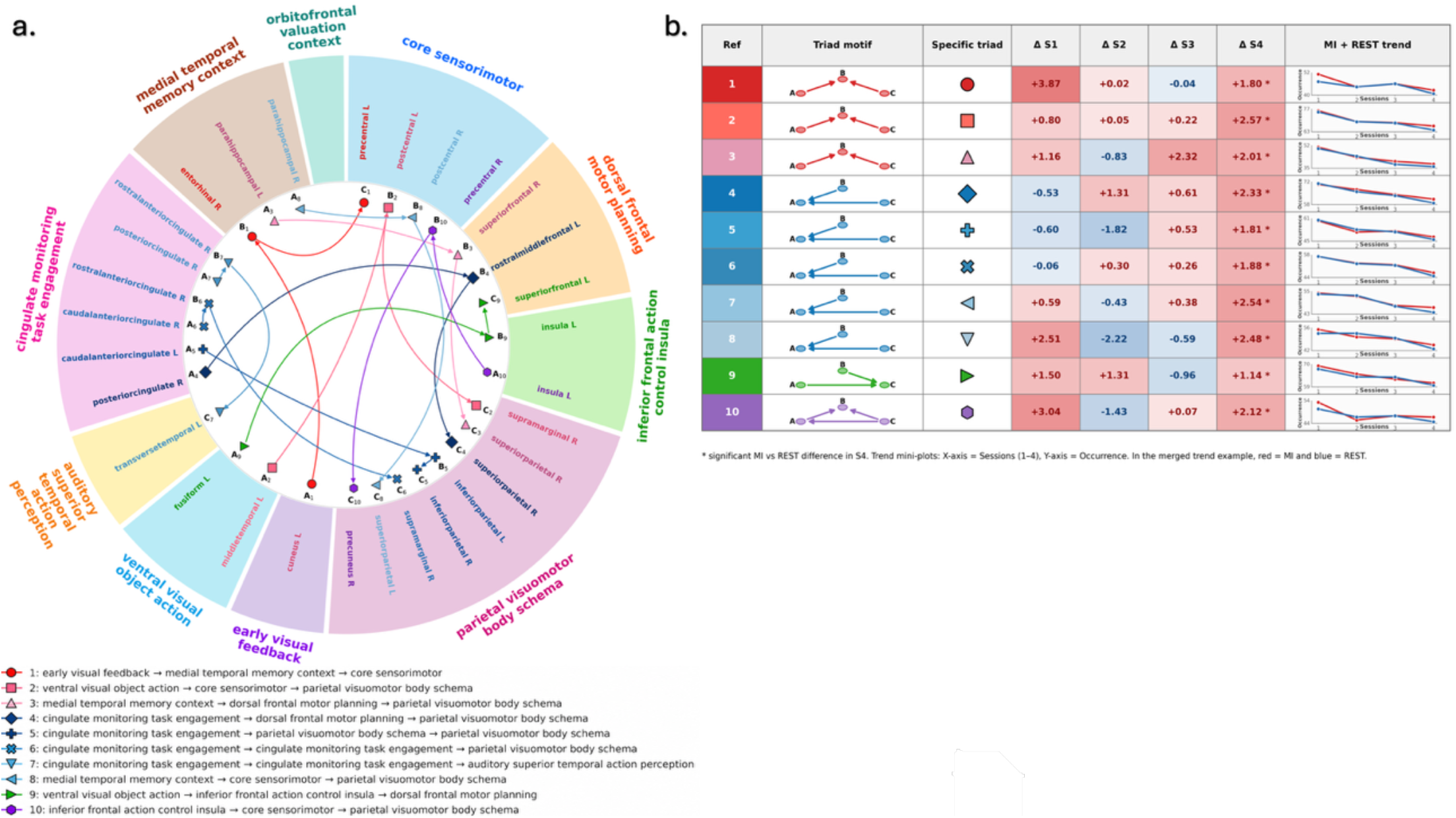
Overview of the ten selected directed triads and their occurrence patterns across training sessions and conditions. (a) Functional connectivity patterns of the ten selected directed triads. Each triad comprises three nodes (A, B, and C), representing distinct functional groups arranged according to their specific directed connectivity patterns. (b) Differences in triad occurrence between MI and REST conditions (Δ = REST − MI) across four training sessions (S1–S4), together with the temporal trends in triad occurrence across sessions for each condition separately. * indicates statistically significant differences between REST and MI.

#### Core-Periphery

To complement the triadic analysis, we examined the core–periphery organization to characterize the relative hierarchical position of functional groups within the broader network. Two main trends emerged from this analysis. First, during motor imagery, ROIs belonging to the core sensorimotor group, particularly the left precentral, left paracentral, and left postcentral regions, tended to occupy more peripheral positions than during resting state (Fig. 5). Second, particularly during the first three sessions, sensorimotor regions showed a progressive shift toward the network periphery as training advanced. In the fourth session, this trend was partially reversed, with a reduction in radial position in both REST and MI. Nevertheless, the relative difference between the two conditions was preserved, with sensorimotor regions remaining more peripheral during MI than during REST (Fig. 5). Session-wise comparisons between REST and MI were assessed using the Wilcoxon test with FDR correction. Significant differences (FDR-corrected p < 0.05) were identified in session 4 for the left paracentral, left postcentral, left precentral, left supramarginal, right banks of the superior temporal sulcus, and right fusiform regions. The only additional significant differences were observed in session 1 for the left postcentral and left precentral regions. When the overall brain dynamics were considered across all training sessions, a significant task effect was identified for the three left sensorimotor ROIs, precentral, paracentral, and postcentral, as well as for the left supramarginal, left posterior cingulate, and right fusiform regions. These regions belong, respectively, to the core sensorimotor, parietal visuomotor body-schema, cingulate monitoring/task-engagement, and ventral visual object-action functional groups. Overall, these functional systems are consistent with the sensory, motor, visuomotor, monitoring, and action-related processes required by the BCI motor-imagery task.

**Fig. 5:**
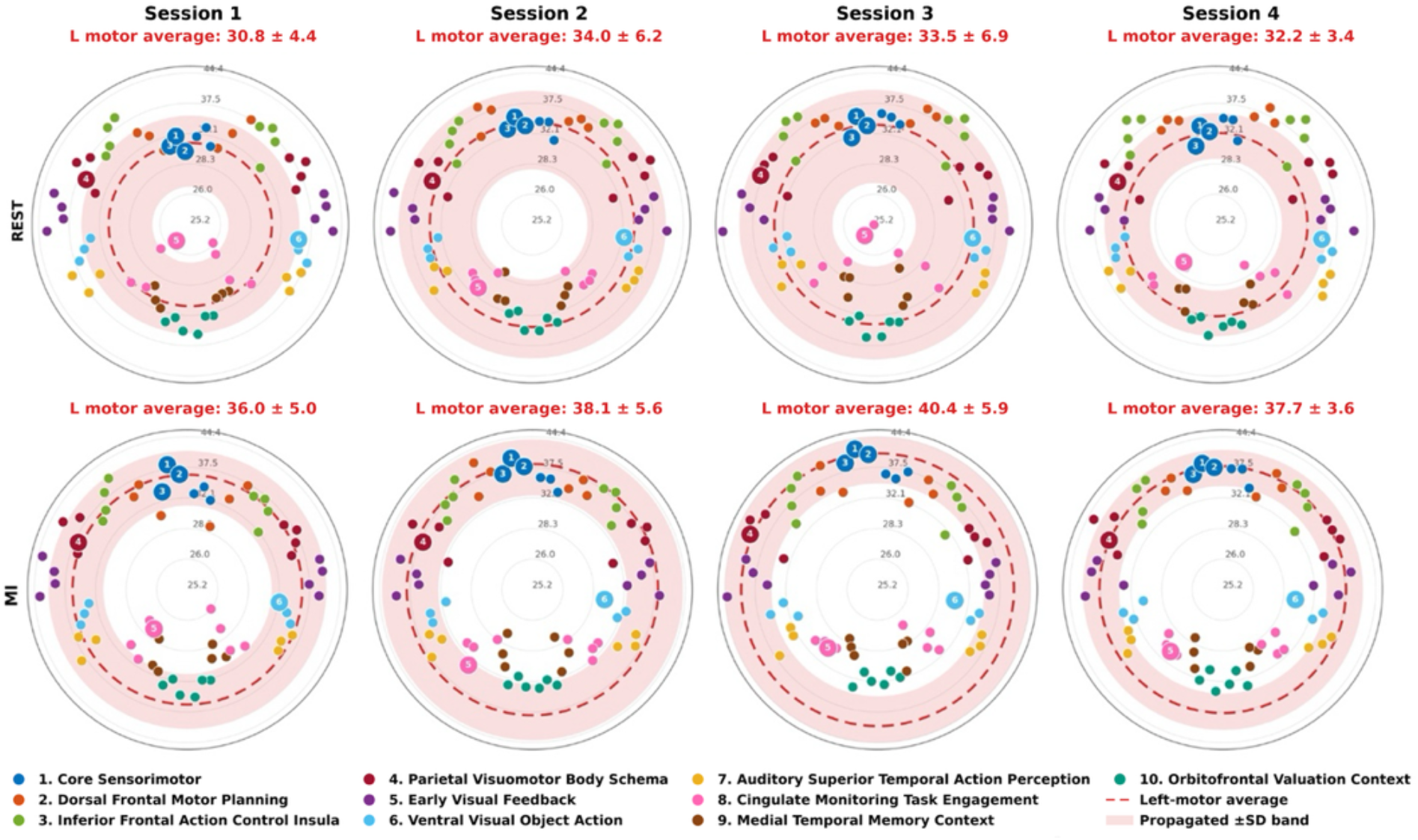
Core–periphery organization across sessions during Rest and motor imagery (MI). The top row shows Rest and the bottom row MI across the four sessions. Each point represents a single ROI, with colors indicating the functional group to which the ROIs belong Fig. 1). ROIs are arranged according to anatomical lateralization, with left-hemisphere regions displayed on the left side of each circle and right-hemisphere regions on the right. The red dashed line and shaded band indicate the mean ± SD of the core–periphery rank for the three left core-sensorimotor ROIs. Larger markers identify ROIs showing a significant task effect in the two-way rank ANOVA: Postcentral L, Precentral L, Supramarginal L, Posterior-cingulate L, Fusiform R

### 3.4 Prediction & Classification Results

The results presented so far support brain-network metrics derived from the W_ij_, and more generally from neuronal avalanche dynamics, as sensitive measures of changes associated with both task and training effects, highlighting their potential as biomarkers. Building on these findings, we extended the analysis to the individual-subject and single-trial level to assess whether the condition-specific PDF observed in the fourth session could be predicted from the trial distributions obtained in the preceding sessions and used for single-trial classification.

Using all ROIs, the classifier achieved a subjects-averaged AUC of 64.5 ± 10.6%. Restricting the analysis to an a priori set of motor-related ROIs identified from the literature did not improve decoding and resulted in the lowest performance (AUC = 62.6 ± 7.1%). In contrast, data-driven spatial selection markedly improved classification. The highest mean AUC was obtained with the a posteriori AUC-based procedure (75.7 ± 11.5%), while PageRank-based ROI selection achieved a comparable AUC of 73.5 ± 7.9%, with no significant difference between the two data-driven approaches (Fig. 6a). The PageRank-derived subset included the left paracentral and precentral cortices, bilateral postcentral, pericalcarine, and inferior parietal regions, and the right lateral occipital cortex (Supplementary Fig. 4a). These findings indicate that graph measures contain substantial information for distinguishing REST from MI but also demonstrate that their decoding potential depends critically on an appropriate definition of the network nodes included in the feature space. Going deeper among the two data-driven approaches, PageRank-based ROI selection provided additional advantages in terms of robustness and computational efficiency. Although its mean AUC was slightly lower than that obtained with the AUC-based strategy, the difference was not significant, and PageRank selection resulted in lower inter-subject variability (Fig. 6a). Moreover, all participants remained above the adopted chance-level[45] threshold of 0.57 when PageRank-selected ROIs were used, whereas below-threshold performance was still observed for some participants with the AUC-based approach. ThePageRank strategy also substantially reduced computational requirements. At the classification stage, the AUC-based procedure required 3.91 ± 0.22 min per subject, compared with 2.40 ± 0.30 min per subject for PageRank-based selection, corresponding to an approximately 39% reduction in computational time, with a significant difference between the two strategies (see Fig. 6b). Although PageRank computation and ANOVA introduced an initial ROI-selection cost around 13.16 s per subject, this overhead was subsequently compensated by the reduced network dimensionality. Specifically, ws-ATM computation decreased from 6.97 min using all ROIs to 1.99 min using the PageRank-selected subset, while the mean cumulative ws-ATM computation across sessions decreased from approximately 17.25 s to 2.76 s per subject. Similarly, computation of the graph measures decreased from 29s across all ROIs to only 966ms after ROI selection.

**Fig. 6:**
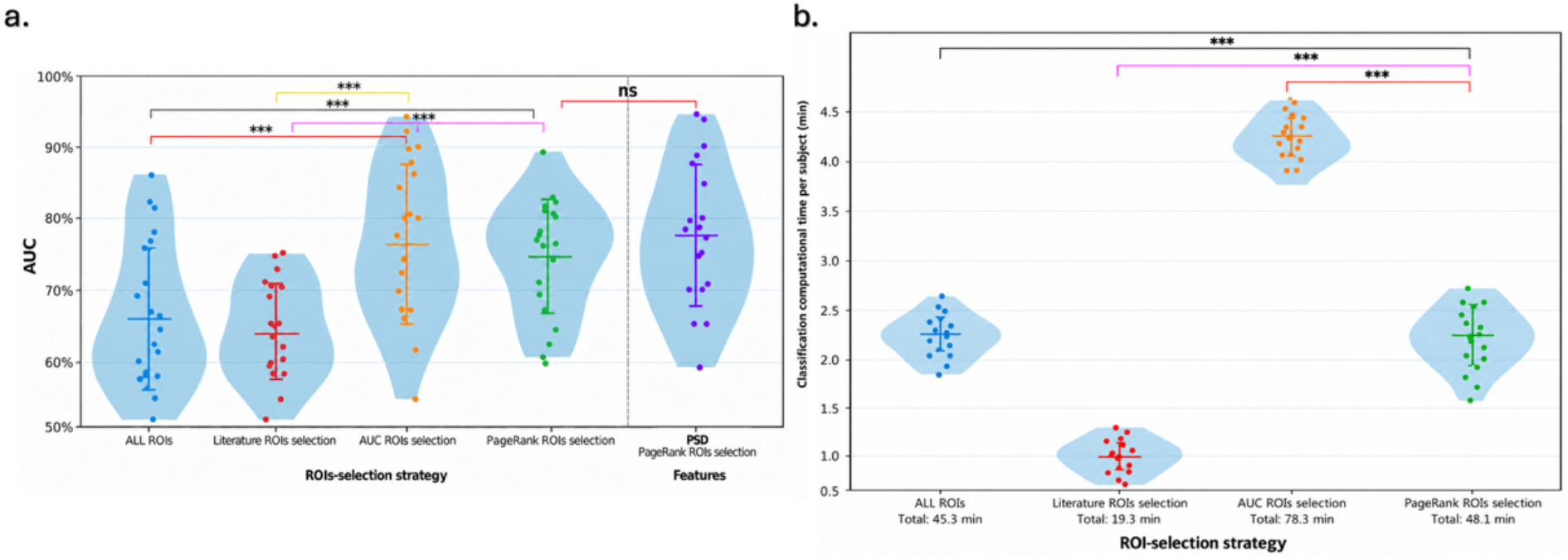
Session 4 classification performance and computational cost across ROI-selection strategies. a) AUC-comparison among different ROIs selection strategies and AUC-comparison between graph-measures and PSD: All ROIs, AUC-based ROI selection, PageRank-based ROI selection, and literature-based ROI selection b) Classification computational time per subject across ROI-selection strategies.

To benchmark graph-derived avalanche features against a conventional EEG representation, we compared their decoding performance with Individual Alpha Frequency (IAF)-based spectral power[46],[47] extracted from the same PageRank-selected ROIs. Spectral features achieved a slightly higher mean AUC than ws-ATM graph measures (0.756 ± 0.121 vs. 0.735 ± 0.079), but the difference was not significant (Fig. 6a). Thus, despite capturing different aspects of brain activity, both representations showed comparable decoding performance, supporting the validity of graph-based features for BCI classification.

## 4 DISCUSSION

A common feature of all analyses in this study is their derivation from directed weighted brain networks. Although less commonly used because of their greater algorithmic complexity, these networks provide a richer description of brain dynamics than undirected representations. Directionality captures the preferential propagation of activity between regions, while edge weights quantify the relative probability of each transition. Together, these properties characterize not only network topology, but also the spatiotemporal organization and preferential flow of neuronal-avalanche propagation across the cortex.

### 4.1 Multi-scale graph measures as biomarkers

Graph-theoretical measures derived from the ws-ATM provide complementary descriptions of neuronal-avalanche propagation across multiple levels of network organization, from the single edge to the whole-network. Edge-wise analysis revealed a clear predominance of higher transition values during REST than during MI, with significant edges mainly directed toward the left sensorimotor system, particularly the postcentral, precentral, and paracentral cortices. This spatial pattern was consistently supported by the nodal measures (Fig. 3), indicating that these regions receive a smaller proportion of incoming transition flow during MI than during REST. The prominent involvement of the contralateral sensorimotor system is consistent with previous MI-connectivity findings, including the reduction in contralateral sensorimotor connectivity and node strength reported in [11]. However, the evolution of these task-related differences across sessions was not monotonic, with highly focal and centered primarily on the left postcentral cortex in Session 1, no significant effects in Session 2 and progressive re-emerge in Session 3 just considering node strength and communicability. This suggests that coordinated changes distributed across multiple connections may modify indirect communication pathways and neighborhood organization even when no single transition is sufficiently strong to reach significance independently.

By Session 4, a broader sensorimotor pattern became evident again, now detectable both at the level of individual transitions and direct node strength, with stronger involvement of the postcentral, precentral, paracentral, and supramarginal regions (Supplementary Fig. 1-3). The early prominence of the postcentral cortex is consistent with evidence that somatosensory plasticity may precede motor-cortical changes during motor learning observed, for example, in [48]. Moreover, in [49] authors reported a decrease in activity in somatosensory area associated with short-term learning while with long-term learning, increases in activity were found in the left primary somatosensory and motor cortex and in the right putamen. Hence, rather than reflecting an absence of plasticity, the reduced direct differentiation observed during the intermediate sessions may therefore represent a transitional stage in which task-related reorganization is first expressed through indirect propagation routes and neighborhood topology before becoming sufficiently consolidated to re-emerge in direct transition strength. The directed nature of the ws-ATM provides an additional level of interpretation by revealing that significant transitions preferentially converge toward left sensorimotor regions (Supplementary Fig. 3), suggesting that these areas act as recipient or convergence sites within neuronal-avalanche propagation. It is in line with previous directed EEG-network analyses proposed by previous studies [50] [51]. In this framework, the observed convergence toward sensorimotor regions may reflect their integrative role in combining body-state, visuospatial, action-planning, and monitoring information during motor imagery.

The triadic analysis (Fig. 4) further supports this interpretation indicating a structured interaction among systems contributing to task monitoring, sensorimotor processing, and visuospatial control. At the same time, the evolution of these motifs across sessions provides evidence of learning-related specialization: although the occurrence of the selected triadic configurations progressively decreased in both REST and MI, their task specificity increased, with all retained motifs showing higher occurrence during MI than REST in Session 4 (Fig. 4). This pattern suggests that training progressively restricts the repertoire of recurrent local propagation patterns while making the remaining configurations increasingly characteristic of the trained MI state. It is consistent with the concept of learning-induced functional autonomy proposed in [52], who showed that motor and visual modules remained consistently recruited during motor-skill learning while their integration progressively decreased, indicating that task-relevant systems become more autonomous with practice. Similarly, in [10] authors reported a progressive reduction in integration among cortical systems specialized in motor imagery and learning, visual attention, and working memory, further supporting the idea that BCI learning is accompanied by increasing functional differentiation among task-related systems. This progressive specialization is also reflected in the core–periphery organization (Fig. 5). The left precentral, postcentral, and paracentral cortices occupied more peripheral positions during MI than during REST, indicating that, despite their central role in the task, they received a smaller proportion of broadly distributed incoming network flow during motor imagery. Across the first three sessions, the left sensorimotor regions progressively shifted toward more peripheral positions during MI, followed by a partial return toward the core in Session 4, while the relative REST–MI difference remained preserved. One possible interpretation is that early and intermediate training progressively promotes greater sensorimotor autonomy, whereas later learning may reflect a second stage of reorganization in which the broader network adapts around a more consolidated MI strategy[53], [54]. This interpretation should nevertheless be considered tentative, particularly given the limited number of training sessions. Core–periphery patterns depend strongly on the analytical framework, including algorithm, modality, and frequency band. Accordingly, differences from previous studies likely reflect methodological sensitivity rather than contradiction [55],. Our longitudinal nodal results further show that training-related reorganization extends beyond sensorimotor regions to frontal, cingulate, parietal, and medial-temporal areas, predominantly during MI, suggesting a progressive redistribution of directed propagation pathways and increasing functional specialization of task-relevant systems. Despite the significant differences observed in specific sensorimotor regions, neither global incoming strength nor global communicability showed a significant overall task effect. Global efficiency, however, showed a complementary pattern, with significantly lower values during MI than REST in Sessions 3 and 4 (Supplementary Fig. 1). This late-emerging reduction is consistent with previous MI-BCI findings [56] and may reflect a transition toward a more segregated network configuration in which the trained MI state relies less on globally efficient shortest-path communication. Across training, global incoming strength and communicability increased in both REST and MI, consistent with previous motor-learning studies reporting increased connectivity strength and degree after practice[9]. Importantly, the increase in global communicability across training is not incompatible with the lower global efficiency observed during MI in the later sessions, because the two measures capture different properties of network communication. Communicability reflects the availability of multiple direct and indirect propagation routes, whereas global efficiency depends specifically on the shortest paths connecting network regions. The coexistence of these effects therefore suggests that training may expand the repertoire and accessibility of possible communication pathways while simultaneously reducing reliance on globally optimized shortest-path integration during MI. In this sense, learning-related plasticity may promote a network that is globally more communicative, yet more functionally segregated and selective during the trained task. This may also indicate that repeated MI training reshapes not only task-related dynamics but the broader baseline network architecture[57].

Taken together, these findings indicate that motor imagery is characterized by a selective reorganization of neuronal-avalanche propagation rather than by a generalized reduction in brain communication. The left sensorimotor system emerges as the main task-related substrate, showing reduced incoming flow, a more peripheral network position, and increasingly specific mesoscale configurations. Across training, coordinated changes in direct and indirect communication, motif recurrence, core–periphery organization, and global integration indicate progressive redistribution and specialization of network dynamics. Because strength, communicability, efficiency, motifs, and core– periphery structure capture complementary aspects of this process, their joint sensitivity to task and longitudinal effects supports ws-ATM-derived graph measures as multiscale biomarkers of brain dynamics.

### 4.2 Multi-scale graph measures as BCI features

#### 4.2.1 Validation Graph measures vs. standard measures

The application of network-based approaches to BCI decoding is a relatively young and unexplored area, yet the existing publications show encouraging results. These works showed that graph measures can provide informative features in motor-imagery classification[12], [58], [59], [60]. These approaches, however, generally derive brain networks from conventional functional-connectivity estimates based on phase synchronization, phase locking, or other statistical dependencies between EEG signals. In contrast, the present framework derives graph measures from a directed-weighted network in which each edge represents the probability that activation of one cortical region is followed by activation of another during neuronal-avalanche propagation. The resulting network therefore describes not only interregional coupling, but the spatiotemporal organization of transient activations propagation across the cortex. By construction, this representation can capture aspects of brain dynamics that are not explicitly encoded by synchronization-based connectivity measures, including the temporal organization of aperiodic neuronal activity.

A second distinctive feature of our approach is its multiscale characterization of network topology. By combining local and global measures, the classifier captures both whole-network reorganization and the regional expression of these changes. This is particularly relevant given the spatial heterogeneity of task- and training-related effects, allowing decoding to exploit both global network dynamics and region-specific alterations that would be lost in purely global summaries. Within this framework, the comparison with IAF-based PSD features provides direct evidence that graph-theoretical summaries of neuronal avalanche dynamics preserve information relevant for BCI decoding. Rather than indicating superiority over conventional spectral features, the results show that ws-ATM-derived graph measures achieve comparable prospective discriminability while capturing a fundamentally different aspect of brain dynamics. This interpretation is consistent with previous analyses of the same dataset, in which features of avalanche dynamics showed competitive within-session performance relative to conventional benchmark methods[16], [17], supporting the discriminative value of neuronal avalanches patterns for separating motor imagery from rest.

Building on this evidence, the present study demonstrates that the same underlying dynamics can be summarized through more compact and interpretable graph-theoretical descriptors, while maintaining useful decoding information under the more demanding condition of cross-session generalization. Moreover, the contribution of the proposed framework extends beyond the choice of network-derived features themselves. Its methodological novelty lies also in using the training-related evolution of these features for prospective decoding, rather than treating inter-session variability solely as a nuisance to be suppressed or corrected.

#### 4.2.2 Longitudinal Classifier Strategy

A central challenge for the practical deployment of motor-imagery BCIs is the non-stationarity of EEG representations across training sessions. Although this variability is commonly treated as a source of decoding instability, our results suggest that part of it may reflect a structured, training-related reorganization that can itself be informative for classification. This is illustrated by the longitudinal evolution of the class-conditional probability density functions (Supplementary Fig. 5): REST and MI distributions of global communicability largely overlap in the early sessions, but progressively diverge, with a clearer separation emerging from Session 3 onward. Importantly, the Session-4 distributions predicted from the preceding sessions closely match the observed distributions while preserving the relative REST–MI separation. This pattern is consistent with longitudinal BCI studies showing that motor-imagery training induces measurable neurophysiological and network-level reorganization and that such changes, also reported in default-mode network[57], can be related to subsequent learning and performance[10], [18]. However, relatively few decoding strategies explicitly exploit the temporal trajectory of training-induced changes to anticipate the neural representations expected in a future session, and, to the best of our knowledge, this trajectory has not yet been directly incorporated into a prospective classification framework. The strong overlap between predicted and observed Session 4 PDFs, together with the preserved REST–MI separation (Supplementary Fig. 5), provides preliminary support for the validity of this strategy. Most subject-specific cross-session transfer-learning methods instead treat inter-session variability primarily as a domain shift to be reduced or compensated for[5], [61], [62]. Stationary spatial-filtering methods, for example, favour features that remain stable across recordings[63], whereas covariance-based and Riemannian approaches reduce session-to-session discrepancies by recentering covariance representations into a common reference space[64]. Similarly, more recent domain-adaptation methods seek session-invariant feature representations by minimizing source–target discrepancies in latent space[65]. Other approaches select or weight historical observations according to their similarity to the current session, as in relevant-session transfer methods[66].

The proposed framework differs from these strategies because it does not primarily seek to suppress or align session-dependent changes, but instead explicitly models their temporal evolution and uses it to predict the class-conditional distributions expected in a subsequent session. A related issue arises in online longitudinal BCI applications, where classifiers are often recalibrated across sessions but can only be updated once data from the new session have become available. In [67], for example, authors investigated upper-limb movement-intention decoding across six sessions while limiting recalibration to promote stable user–decoder interaction. The classifier from the previous session was retained when performance remained sufficiently stable, but when performance deteriorated, a new model was trained using calibration data collected in the current session. Thus, even in this adaptive setting, recalibration depended on observations from the target session. In contrast, the predictive nature of the present framework could, in principle, support pre-session recalibration by estimating the expected REST and MI distributions from the individual’s previous training trajectory before the new session begins. Such an approach could reduce the time patients spend in the laboratory at the beginning of subsequent BCI sessions while also enhancing user autonomy and facilitating the use of this technology in less controlled environments. However, its effectiveness in a truly online setting remains to be prospectively validated.

#### 4.2.3 Advantage of ROIs selection

To reduce calibration and computational burden after initial BCI training, we assessed whether spatial feature reduction could preserve meaningful information while simplifying decoding. Feature selection is a key component of EEG-based BCI pipelines, as it can reduce model complexity, computational cost, and calibration time[5], [68]. Conventional ROI/channel selection can be broadly divided into *a priori* and *a posteriori* approaches. *A priori* strategies restrict the analysis to anatomically or neurophysiologically defined sensorimotor regions, offering simplicity and interpretability but potentially overlooking the broader network involved in MI. Indeed, meta-analyses show that MI recruits premotor, parietal, somatosensory, and associative regions, with primary motor cortex involvement being less consistent[38], [39]. In contrast, *a posteriori* approaches select ROIs according to decoding performance, for example by ranking features or iteratively adding or removing channels[69] but repeated optimization on classification accuracy can introduce selection bias and inflate generalization estimates[70].

We therefore introduced a complementary data-driven strategy independent of classification performance, selecting ROIs from the Condition × Session interaction of PageRank centrality. This identifies regions whose network relevance captures both REST–MI differences and their evolution across training, thereby incorporating the longitudinal dimension of BCI learning into spatial feature selection[71]. Importantly, PageRank-based ROI selection for prospective classification used only the first three sessions, with Session 4 kept completely unseen until final testing, preventing test-session information from influencing spatial selection. Although its mean decoding performance was comparable to the AUC-based approach and significantly better than the literature-based selection, our method yielded lower inter-subject variability and reduced computational time by about 39% (3.91 ± 0.22 to 2.40 ± 0.30 min per subject), with additional savings in ws-ATM construction and graph-metric computation. These findings suggest that the main advantage of PageRank-based selection is not necessarily a direct increase in classification accuracy, but rather a favorable compromise between neurophysiological interpretability, sensitivity to training-related reorganization, robustness across participants, and computational efficiency.

With the same emphasis on neurophysiological relevance, we performed an additional exploratory analysis to assess the information retained after dimensionality reduction by reconstructing the ws-ATM using the selected ROI subset identified across all four sessions. At the local scale, the reduced network reproduced the main patterns observed using the complete set of ROIs, while at the global scale significant REST–MI differences emerged in Session 4 for both global communicability and global node strength (Supplementary Fig. 4b). Together, these observations indicate that restricting the network to regions exhibiting task-dependent longitudinal modulation can retain, and in some cases enhance, the expression of relevant large-scale network differences, supporting the potential of this ROI-selection strategy for computationally efficient longitudinal BCI applications.

### 4.3 Limitation and future prospective

The proposed graph-based approach provides an alternative to conventional PSD features for Motor-Imagery decoding. Although it did not yield a consistent improvement in decoding performance in the present study, it offers a complementary perspective on brain network dynamics. Similarly, PageRank-based ROI selection represents an alternative to AUC-based feature selection, although no additional performance gains were observed in the current experimental setting. Future work should therefore investigate hybrid feature representations rather than replacing conventional features with network measures. Previous studies support this direction: results in [11] showed that combining connectivity-derived node strength with PSD improved classification in most subjects, while in [72] authors reported gains when graph measures were combined with CSP-based power features, despite CSP performing better when used alone. These findings suggest that network and spectral features capture complementary information. A promising extension would therefore be to combine ws-ATM graph metrics with PSD features within the PageRank-selected ROI subset, potentially improving decoding while preserving dimensionality reduction.

A second limitation concerns the limited number of recording sessions, which prevents us from conclusively validating the hypothesis that the core–periphery reorganization observed in Session 4 reflects an adaptation process induced by training. Future studies should therefore investigate this hypothesis using datasets with a larger number of longitudinal training sessions. Increasing the number of sessions would also provide a more reliable characterization of the distributions associated with the two experimental conditions and could consequently improve the robustness and accuracy of the classification model.

More generally, the framework proposed in this work is intended to evolve toward an online BCI implementation. In this perspective, the initial sessions could be used as a calibration phase to optimize the population-level model and identify a fixed set of informative features and relevant ROIs or EEG channels. Once these parameters are established, subsequent sessions could be processed in an online setting in which only single-trial classification is required. In the present study, this scenario is approximated by using Session 4 as an independent test set. Importantly, with PageRank-based selection, single-trial classification required around 47.25 ± 3.45 ms, well below typical real-time BCI latency requirements around 100 ms for prosthetic applications. These preliminary computational results therefore support the feasibility of extending the proposed framework toward a real-time online BCI implementation.

## 5. CONCLUSION

In this work, we introduced a BCI framework integrating neuronal-avalanche dynamics, graph theory, longitudinal modelling, and data-driven ROI selection. Unlike approaches focused primarily on classification performance or computational efficiency, our framework places neurophysiological interpretability at its core by first assessing whether ws-ATM-derived graph measures provide meaningful biomarkers of motor imagery and training-related brain reorganization across multiple topological scales. We also introduced a different perspective on inter-session variability. Instead of considering BCI non-stationarity exclusively as a source of variability to be removed or compensated for, we explicitly modelled the evolution of feature distributions across training sessions and used this information to predict brain states in a subsequent, unseen session. Finally, the PageRank-based ROI-selection strategy exploited the interaction between task and training to identify a reduced set of neurophysiologically informative regions, providing a compromise between interpretability, prospective decoding performance, inter-subject robustness, and computational efficiency. Together, these components aim to shift part of the feature-selection and computational burden to an initial offline calibration phase, simplifying subsequent BCI use. An initial set of supervised sessions could identify subject-specific features and brain regions, after which the same configuration could be applied more autonomously during later rehabilitation or training. Reducing calibration demands and expert supervision may facilitate more frequent use in physiotherapist-supervised or home-based settings, supporting future neurorehabilitation applications.

