## Supplementary Materials for "Brain Network Dynamics from Neuronal Avalanches for Longitudinal Motor-Imagery Decoding"

| Functional group | Desikan–Killiany ROIs | Reference |
|---|---|---|
| **1. Core sensorimotor** | Precentral; Postcentral; Paracentral | Hill et al. (2019) |
| **2. Dorsal frontal motor planning** | Superior frontal; Caudal middle frontal; Rostral middle frontal | Wise et al. (1997) |
| **3. Inferior frontal action control / insula** | Pars opercularis; Pars triangularis; Pars orbitalis; Insula | Aron et al. (2014) |
| **4. Parietal visuomotor / body-schema** | Superior parietal; Inferior parietal; Supramarginal; Precuneus | Andersen & Buneo (2002) |
| **5. Early visual feedback** | Pericalcarine; Cuneus; Lingual; Lateral occipital | Roelfsema & de Lange (2016) |
| **6. Ventral visual object / action processing** | Fusiform; Inferior temporal; Middle temporal | Wurm & Caramazza (2022) |
| **7. Auditory / superior temporal action perception** | Superior temporal; Transverse temporal; Banks of the superior temporal sulcus | Alho et al. (2014) |
| **8. Cingulate monitoring / task engagement** | Caudal anterior cingulate; Rostral anterior cingulate; Posterior cingulate; Isthmus cingulate | Botvinick et al. (2004) |
| **9. Medial temporal memory / context** | Entorhinal; Parahippocampal; Temporal pole | Eichenbaum et al. (2012) |
| **10. Orbitofrontal valuation / context** | Lateral orbitofrontal; Medial orbitofrontal; Frontal pole | Knudsen & Wallis (2022) |

**Note.** Left- and right-hemisphere homologues were assigned to the same functional group.

***Supplementary Table 1. Functional grouping of cortical regions of interest (ROIs) based on the Desikan–Killiany parcellation.*** *Cortical ROIs were grouped post hoc into ten functionally defined categories based on their anatomical locations and functional roles. Each functional group comprises anatomically and functionally related ROIs, with assignments supported by the corresponding literature references. Homologous ROIs from the left and right hemispheres were assigned to the same functional group.*

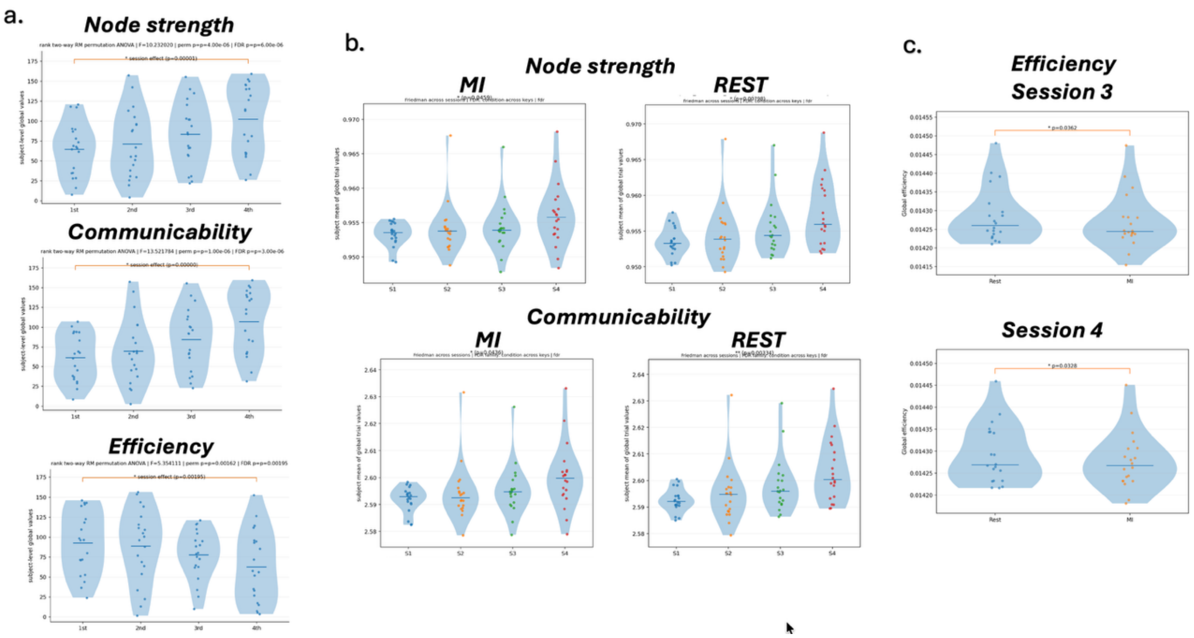


***Supplementary Fig. 1:* *Global network measures across sessions and conditions.*** ***a)*** *Session effect on node strength, communicability, and efficiency assessed using a two-way ANOVA.* ***b)*** *Session-related changes evaluated separately within the REST and motor imagery (MI) conditions using Friedman tests, with FDR correction for multiple comparisons.* ***c)*** *Task-related differences between REST and MI within each session assessed using Wilcoxon signed-rank tests, with FDR correction.*

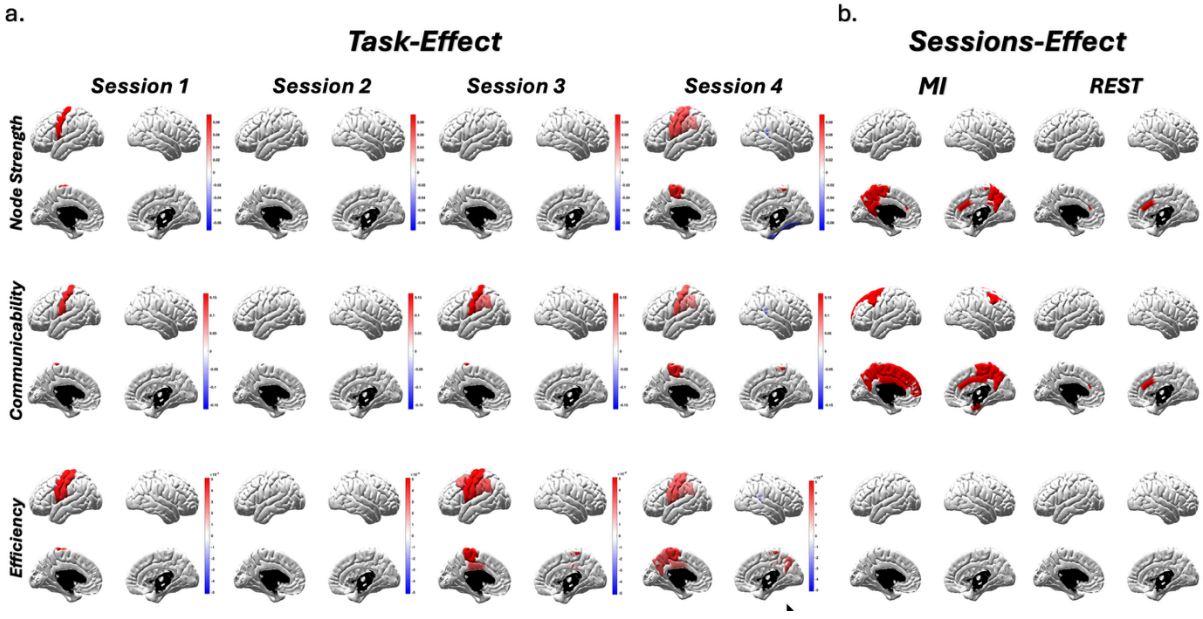


**Supplementary Fig. 2. Spatial distribution of significant task and session effects across network metrics.** Brain regions exhibiting statistically significant differences in node strength, communicability, and efficiency, evaluated separately for task and session effects. **(a) Task effect:** Spatial distribution of regions of interest (ROIs) showing significant differences between motor imagery (MI) and resting-state (REST) conditions within each training session (S1–S4), independently assessed using t-tests with false discovery rate (FDR) correction ($p < 0.05$). Red and blue indicate positive and negative differences, respectively, with color intensity reflecting the magnitude of the difference. **(b) Session effect:** Spatial distribution of ROIs exhibiting significant changes across the four training sessions, analyzed separately for MI and REST conditions. Red highlights ROIs showing significant session-related effects.

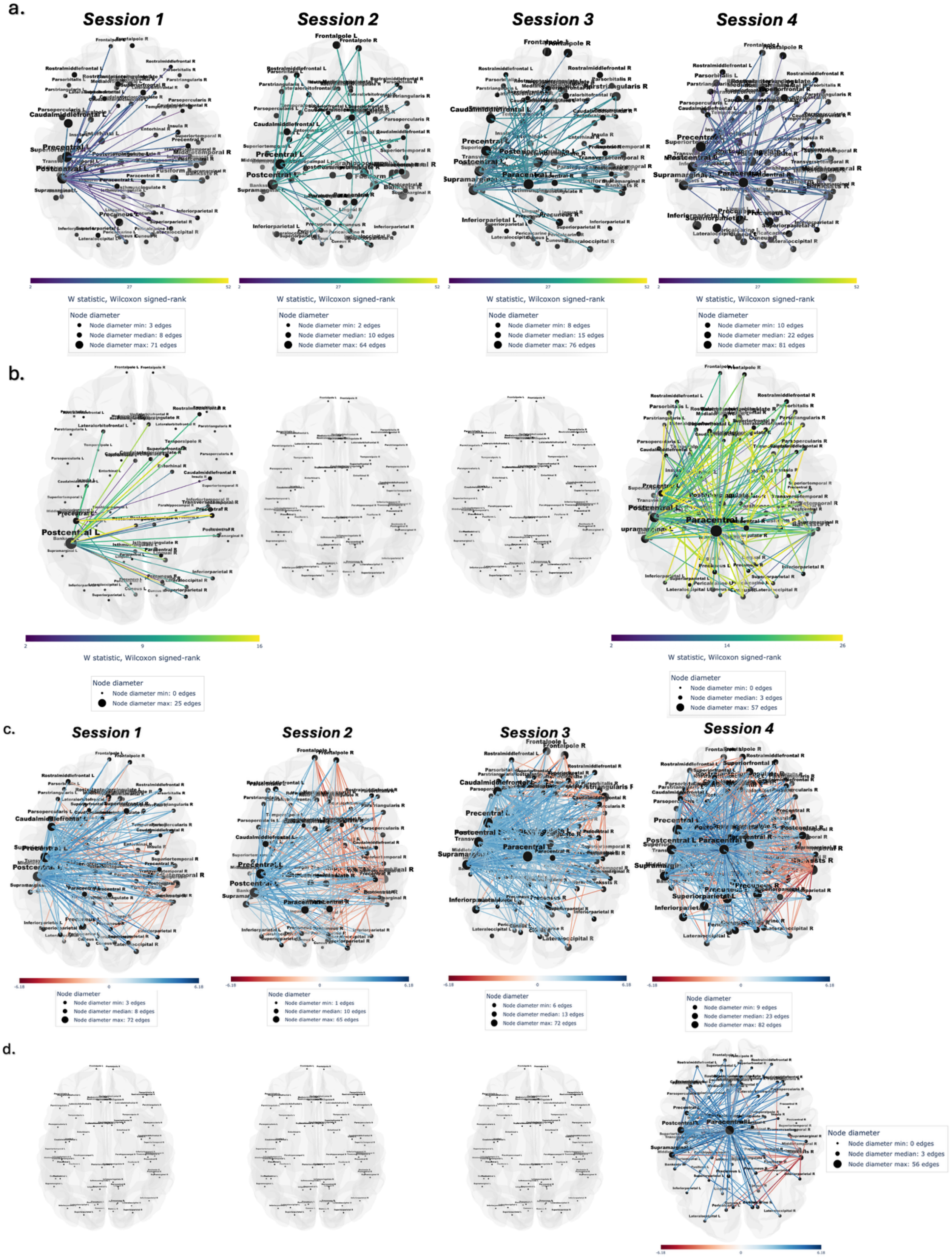


***Supplementary Fig. 3: Edge-wise differences in functional connectivity between motor imagery (MI) and resting-state (REST) conditions across training sessions.*** *Spatial representation of significant edge-wise differences between REST and MI within each of the four training sessions (S1–S4), assessed using Wilcoxon signed-rank tests and paired t-tests.* ***a, b) Wilcoxon signed-rank test:*** *Significant connections identified before (a) and after (b) false discovery rate (FDR) correction for multiple comparisons. Edge colours represent the Wilcoxon signed-*

*rank statistic (W), as indicated by the corresponding colour scales.* ***c, d) Paired t-test:*** *Significant connections identified before (c) and after (d) FDR correction. Edge colors indicate the direction and magnitude of the t-statistic (REST − MI), with blue representing significantly greater connectivity during REST than MI (positive t-values) and red representing significantly greater connectivity during MI than REST (negative t-values). Across all panels, node size is proportional to node degree, defined as the number of significant connections involving each region of interest (ROI). Only connections meeting the statistical significance threshold (*$p < 0.05$*, uncorrected or FDR-corrected, as appropriate) are displayed.*

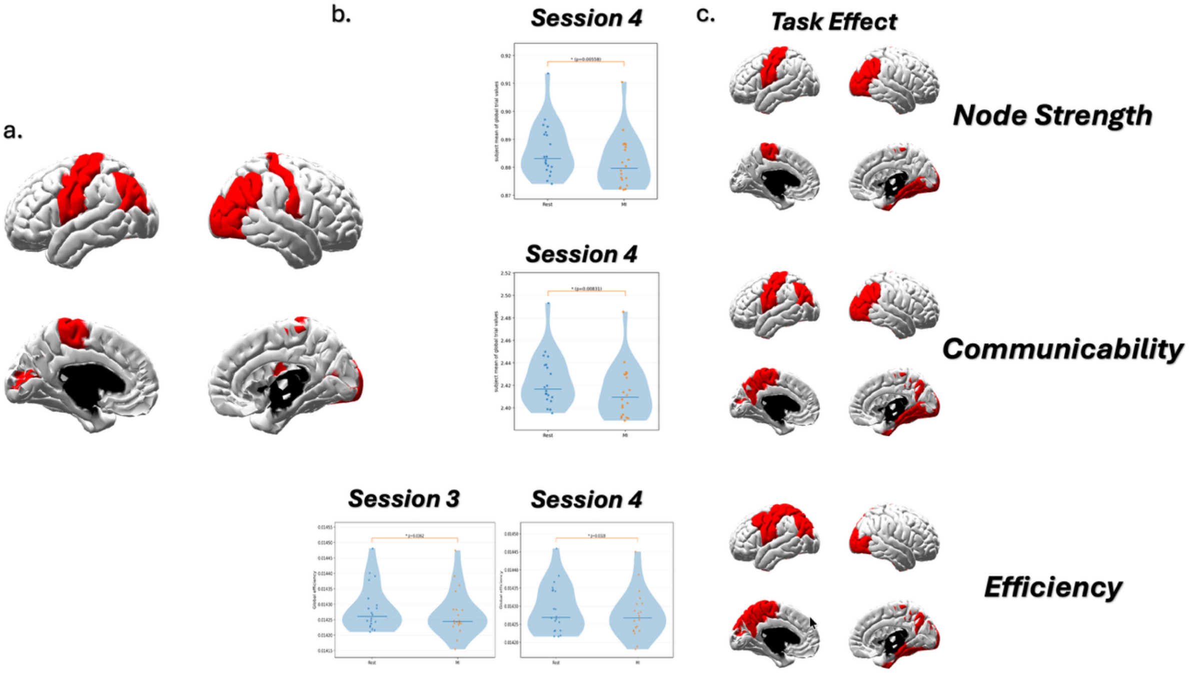


***Supplementary Fig. 4. Region of interest (ROI) selection and significant task-related differences in global and local network metrics. a) PageRank-based ROI selection:*** *Spatial distribution of the nine ROIs selected using PageRank-based feature selection across the four training sessions (S1–S4), highlighted in red.* ***b) Global network metrics:*** *Violin plots illustrating significant differences between resting-state (REST) and motor imagery (MI) conditions in node strength and communicability during session S4, and efficiency during sessions S3 and S4, as identified by Wilcoxon signed-rank tests following false discovery rate (FDR) correction (*$p < 0.05$*). Individual observations and summary statistics are overlaid on the distributions, with the corresponding significance levels indicated above each comparison.* ***c) Local network metrics:*** *Spatial distribution of ROIs exhibiting significant main effects of task (MI vs. REST) across all four training sessions for node strength, communicability, and efficiency, as identified by two-way ANOVA with FDR correction (*$p < 0.05$*). Red regions indicate statistically significant task-related effects, whereas gray regions indicate non-significant ROIs.*

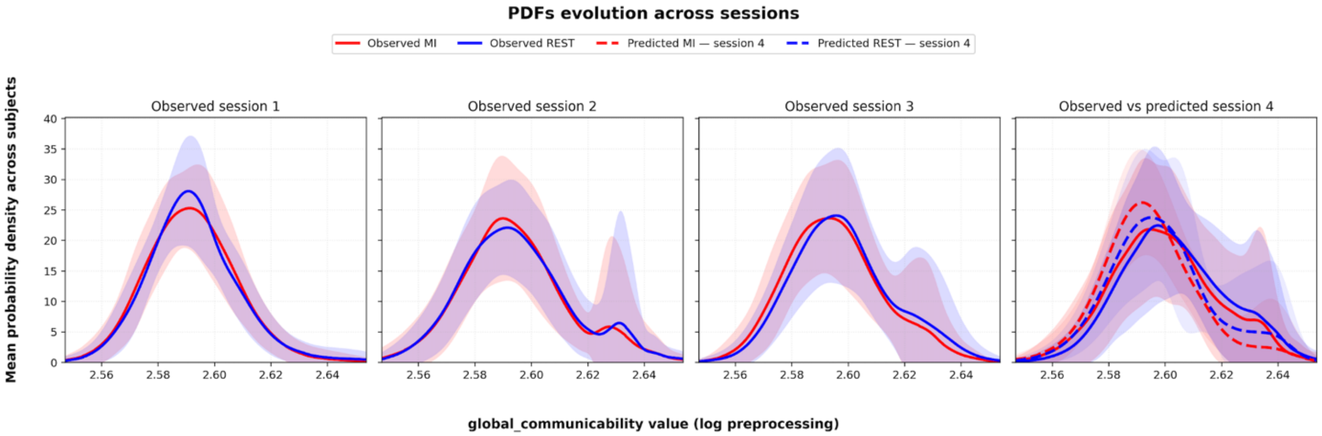


***Supplementary Fig. 5: Evolution of the subject-averaged probability density functions (PDFs) of Global Communicability across sessions.*** *Solid blue and red lines represent the mean observed PDFs across subjects for the REST and motor imagery (MI) conditions, respectively, in Sessions 1–4. The corresponding shaded areas indicate the inter-subject variability (mean ± SD). In Session 4, dashed blue and red lines show the predicted PDFs for REST and MI, respectively, obtained from the model based on the preceding sessions.*
